# Mice and humans evaluate odor stimulus strength using common psychophysical principles

**DOI:** 10.1101/2025.08.16.669760

**Authors:** Beatrice Barra, Robert Pellegrino, Jacqueline Zhao, Christiane Danilo, Aiden Streleckis, Timothy Reizis, Dmitry Rinberg, Joel D. Mainland

## Abstract

Sensory systems translate physical stimuli from the environment—such as light, sound, or chemicals—into signals that the brain can interpret. Across these systems, the amplitude of a stimulus is represented by its perceived intensity. Although previous research has extensively studied how the brain represents physical stimuli, less is known about how it represents perceptual variables such as stimulus intensity. This is primarily due to the difficulty in measuring perceptual responses in animal models, where neural recordings are more accessible. In this study, we use mouse olfaction as a model system to develop a framework for measuring perceived odor intensity. We begin by employing a two-odor concentration classification task to demonstrate that both mice and humans assess stimulus amplitude using a common perceptual scale. We then show that this scale corresponds to intensity. Finally, we apply this method to determine isointense concentrations of different odorants in mice. Our approach offers a powerful tool for testing hypotheses about the neural mechanisms underlying perceived odor intensity, potentially enhancing our understanding of olfactory processing and its neural substrates.

## Introduction

Humans can easily compare the loudness of a piano to that of a violin, even though these instruments produce markedly different sound spectra. Similarly, we can judge the brightness of different colors or the strength of different odors. Perceived intensity is a fundamental property of sensation that spans all sensory stimuli and describes their magnitude. Judging stimulus intensity is critical across all sensory systems and can impact the attention devoted to a stimulus (i.e., a loud sound can prompt a fast reaction). Conversely, abnormal intensity perception can signal sensory dysfunction and is associated with neurological and psychiatric conditions.

Early psychophysicists such as Fechner and Stevens sought to use measures of human sensation to infer the internal mechanisms of intensity perception (Stevens 1957; Fechner 1948). Their work led to general laws describing intensity perception across sensory modalities. Subsequent studies expanded on these foundations, and characterized differences in intensity perception across different stimuli within each sensory system (Gescheider 1997). In all sensory systems, *perceived intensity* increases monotonically with *stimulus amplitude*, however the specific relationships between these variables are stimulus dependent. For example, in audition, the relationship between loudness and sound pressure amplitude depends on the sound’s frequency (Moore 2012). In olfaction, perceived intensity scales with odorant concentration in an odorant-specific manner (Chastrette et al. 1998; Cain 1969). Although significant progress has been made in linking physical stimulus variables to neural responses (Bolding and Franks 2017; Spors et al. 2006; Schreiner and Malone 2015; Phillips 1987; Heinz et al. 2005; Micheyl et al. 2013; Bensmaia 2008; Muniak et al. 2007; James et al. 2019; Dyballa et al. 2024), the neural basis of intensity perception remains poorly understood. This gap reflects both the difficulty of measuring perceptual and neural responses in the same animal and the lack of precise mappings between physical and perceptual variables. Establishing isointense stimuli would provide a critical tool for disambiguating perceptual strategies, such as distinguishing whether animals rely on stimulus identity or intensity.

In olfaction, the work in rodents has identified several neural correlates of odor concentration in the olfactory bulb and piriform cortex (Mainland et al. 2014), including single unit spike rates, spike latencies (Spors et al. 2006; Sirotin et al. 2015), spatial extent of responses (Wachowiak and Cohen 2001), and ensemble synchrony (Bolding and Franks 2017). However, to identify which of these signals relate to perceived odor intensity, it is necessary to dissociate perceived intensity from stimulus concentration. Currently, this dissociation is only achievable in humans, where comprehensive multi-area neural recordings are impractical. Accurately measuring isointense concentrations in a mouse model would finally enable mechanistic studies on the neural coding of perceived odor intensity, and aid in a broader understanding of odor coding.

Moreover, it would shed light on whether animals and humans adopt similar psychophysical principles to evaluate the intensity of sensory stimuli.

Here, we adapted a two-odor concentration classification (2OCC) task that suggested that intensity disparities across odors led to systematic errors in a concentration classification task (Wojcik and Sirotin 2014). We implemented this task in both mice and humans and found that both species evaluate odor concentration using a common perceptual scale, which we demonstrate corresponds to perceived intensity. We then applied this paradigm in mice to estimate isointense concentrations for four odorants at three different intensities. These estimates allowed us to establish the odor-specific relationships between perceived intensity and odorant concentration in mice. Our approach enables the use of mice to investigate the neural basis of intensity perception and, more broadly, the principles of odor coding.

## Results

### Mice discriminate concentrations across odors using a common scale

To measure perceived intensity in mice, we adapted a 2OCC paradigm previously developed in rats (Wojcik and Sirotin 2014). First, head fixed mice (n=4) were trained to discriminate four high concentrations from four low concentrations of a single odor (ethyl tiglate, ET), by licking left or right waterspouts (**Figure 1A, B**). Concentrations were distributed uniformly on a logarithmic scale spanning one order of magnitude. Odor stimuli at specific concentrations were delivered using an air dilution olfactometer (**Figure S1**). Mice learned to perform the task over weeks and training was terminated upon reaching a 75% performance criterion and a decision boundary estimated to be ±0.071 from the middle of logarithmic concentration range (**Figure S1D**). Once trained on a reference odor, they were able to quickly generalize to different odorants, namely ethyl butyrate (EB) and 2-heptanone (2Hep) (**Figure 1B)** and upon the first presentation of the new odorants, displayed performances were not different from performances with ET (Friedman chi-squared test, p = 0.779, **Figure 1C**).

**Figure 1.**
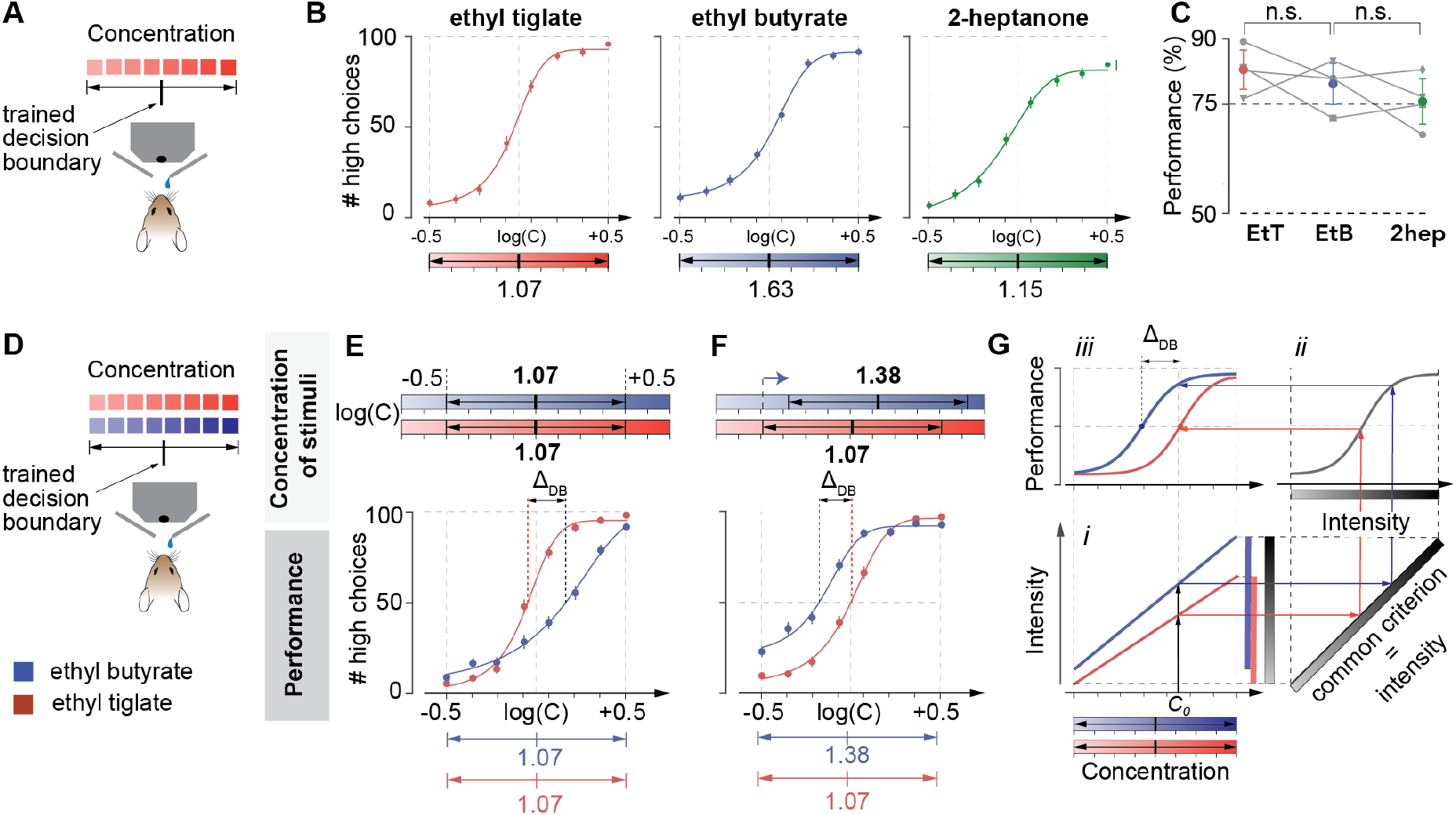
Mice classify odorant concentrations using a common scale. **A**. Task design. Mice (n=4) were trained to categorize eight concentrations of a single odorant by licking the left/right ports for the four lowest/highest concentrations. Concentrations were distributed uniformly on a logarithmic scale, spanning one order of magnitude. Correct responses were rewarded with a drop of water. **B**. Performance at the final training session with ethyl tiglate (ET, red) and the first session with novel odorants ethyl butyrate (EB, blue) and 2-heptanone (2H, green). The x-axis is centered on the logarithm of odorant concentration in parts per million (ppm) and extends +0.5 units in logarithmic space. **C**. Overall performance across all three odorants. Data points show means ± SD across mice. Gray lines and symbols correspond to performance of individual mice. No significant differences were found across odorants (Friedman chi-squared test, p = 0.779). **D**. Two-odorant task design. Mice were rewarded for licking the left/right ports for the four lowest/highest concentrations of both odorants. **E**. Mean psychometric curves (n = 4) for the two-odorant task with ET (red) and EB (blue) using the same concentration ranges (midpoint concentration range for both odorants is 10^1.07^ ppm). Data points show means ± SEM. **F**. The same as E, but with a shifted concentration range for EB. **G**. Schematic of behavioral task performance under a common intensity criterion. If the concentration ranges of the two odorants correspond to different perceived intensity ranges (i), applying a common criterion (ii) would yield performance differences determined by relative differences in intensity perception between stimuli (iii). Δ_DB_ indicates the relative difference between decision boundaries for individual odorants.

The 2OCC paradigm relies on the observation that when stimuli from two different odorants (8 concentrations of each odorant, for a total of 16 possible stimuli) are presented in the same experimental session, animals may display differences in their odor-specific performance (Wojcik and Sirotin 2014). When ET and EB stimuli were interleaved in the same session (**Figure 1D)**, performance differed by odorant identity. ET stimuli were more often misclassified as “high”, despite being rewarded as “low”, while EB stimuli were more often misclassified as “low”, despite being rewarded as “high stimuli” (**Figure 1E)**. Shifting the concentration range of EB systematically altered these biases: at higher EB concentrations, mice classified EB stimuli as more “high-like” than ET (**Figure 1F)**.

These odor-specific biases suggest that mice do not use independent decision boundaries for each odorant. Instead, they appear to rely on a shared perceptual scale, consistent with the notion that odorants are judged according to their perceived intensity (Wojcik and Sirotin 2014). To understand how this would produce the odor-specific performances observed experimentally, we can analyze a scenario where for some concentration range odorant A evokes higher intensity perception than odorant B (**Figure 1Gi)**. A mismatch of the intensity curves would lead to an overall larger range of intensities (see gray bar), compared to the intensity ranges for individual odorants. If the animal is relying on this internal variable to execute the task, the performance curve will look like a sigmoidal function of intensity, shown at **Figure 1Gii**. This in turn will lead to a different level of performance for two odorants at the same concentration, *C*_*0*_. Therefore, the resulting psychometric curves for individual odorants in concentration coordinates will be shifted relative to each other by some interval, Δ_DB_ (**Figure 1Giii**). Changing the range of concentrations of individual odorants will affect this shift between the psychometric curves.

In summary, our results support the hypothesis that mice evaluate odorant concentrations using a common perceptual variable that is shared across odorants and we assume that this variable is the perceived odor intensity.

### The common perceptual scale used to evaluate odorant concentration is odor intensity

To test whether this common scale corresponds to perceived intensity, we performed analogous experiments in humans, where subjective intensity can be measured directly. A panel of participants (n=17) performed the 2OCC task with two odorants, acetal (AC) and 2-heptanone (2H) and, similarly to the mouse, received feedback (correct/incorrect) on every trial. The panel also rated subjective odor intensity across a wide range of concentrations in a separate session (**Figure 2**). AC concentrations were fixed across sessions, while 2H concentrations were systematically varied. As in mice, human psychometric functions shifted relative to the rewarded boundary. **Figure 2A** shows the average behavioral performance of n = 17 subjects: panelists exhibited an increased number of *high* choices for 2H compared to ET, despite 2H being delivered at lower concentrations with respect to AC. This result underscores that mice and humans evaluate odorant stimuli using a similar psychophysical strategy, based on a common perceptual scale. Moreover, the *increased rate* of high choices for stimuli at *lower concentrations*, reinforces the notion that similar concentrations of different odorants can be perceived as extremely different, depending on each specific odorant. Next, we reasoned that, like in the mouse, a decrease in 2H concentrations would attenuate the difference in performance for the two odorants. Indeed, a moderate decrease in the concentration range of 2H performance curves reflected a similar number of *low* choices for 2H compared to AC (**Figure 2B**), and a further decrease in 2H concentrations induced an increased number of *low* choices for 2H compared to AC (**Figure 2C**).

**Figure 2.**
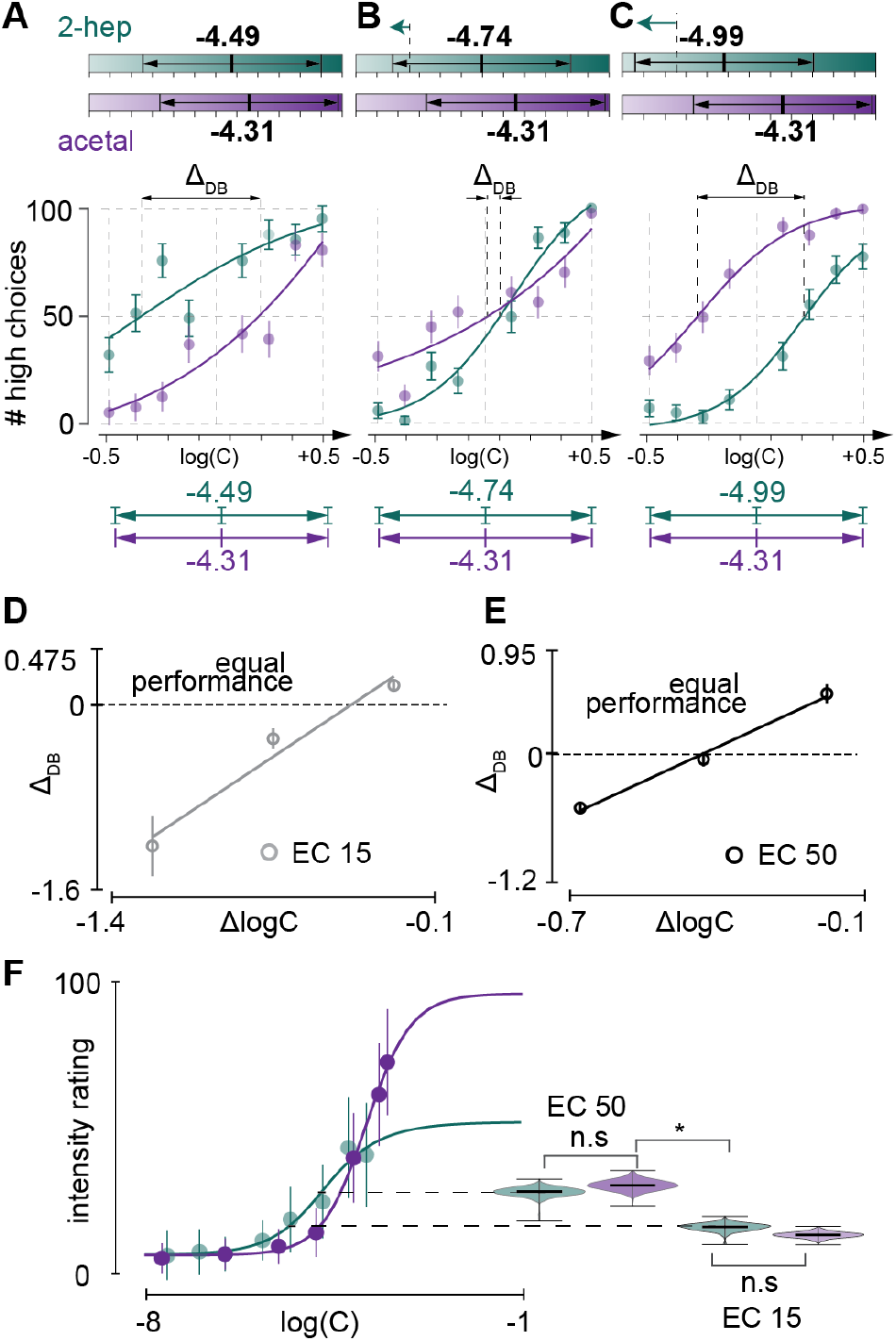
Humans also apply a shared intensity criterion across odorants. **A**. Two-odorant classification performance in human participants (n=17) for a fixed set of eight acetal (AC) concentrations and eight 2-heptanone (2H) concentrations. The midpoints of the concentration ranges were -4.49 (2H) and -4.31 (AC) where values represent the logarithm of odorant concentration (vol/vol). Data points show means ± SEM. **B**. Same as A, with the 2H concentration range centered at -4.74. **C**. Same as A and B, with the 2H concentration range centered at -4.99. **D**. Performance mismatch index (Δ_DB_) plotted against the difference between 2H and AC concentration ranges. The 2H range was centered at 15% (EC15) of its maximum perceived intensity. **E**. Same as D, but with the 2H concentration range centered at 50% (EC50) of its maximum perceived intensity. **F**. Perceived intensity as a function of concentration. Data points show means ± SD. Violin plots show the mean perceived intensities of concentration pairs that yielded equivalent performance (Δ_DB_ =0) at the EC15 and EC50 concentration ranges. n.s. indicates p > 0.05, ^*^ indicates p < 0.05 (two-sided bootstrap resampling).

If the common scale is indeed intensity, then psychometric curves should overlap when isointense concentrations are presented. In other words, if the relative shift of psychophysics concentration discrimination curves for individual odorants, Δ_DB_, is proportional to the intensity mismatch across odorants, then for isointense odorant concentrations the shift should be zero: Δ_*DB*_= 0. We measured the difference in performance Δ_DB_ as the difference between the category boundary values for the two odors. We define the point of subjective equivalence (PSE) as the concentration where Δ_DB_ = 0 and a subject treats the two odors as equally intense. We plotted the distance on the common scale against the difference in concentration for the three sessions and used a linear fit to identify the PSE (**Figure 2D)**. We repeated this procedure at two different concentration ranges, that corresponded approximately to the 15% (EC15, **Figure 2D**) and 50% (EC50, **Figure 2E**) of the dynamic range of intensity of 2H to predict isointense concentrations at two different intensities ranges.

Independent ratings using the general Labeled Magnitude Scale (Green et al. 1996; Bartoshuk et al. 2003) (gLMS) confirmed these predictions. Participants rated the intensity of ten concentrations of AC and 2H (**Figure 2F**). The two pairs of isointense concentrations predicted by the 2OCC were explicitly rated as isointense (**Figure 2F, right**). For comparison, concentrations associated to different intensity ranges resulted in significantly different intensity ratings. The relationship between Δ_DB_ and 2H concentration changes was also steeper at the higher intensity. We conclude that humans evaluate odorant concentration on a common intensity scale and hence the 2OCC task is an effective tool to measure isointense concentrations of odorants. We then applied this paradigm to measure isointense concentrations in a mouse.

### Isointense concentrations in mice are transitive across odors

We performed the 2OCC task in mice at three intensity ranges with four odorants: ethyl tiglate ET), ethyl butyrate (EB), 2-heptanone (2H) and acetophenone (ACE). We then reasoned that, for the 2OCC task to accurately estimate isointense concentrations of the different odorants, isointense concentrations across odorants should be transitive, meaning that matched concentrations do not depend on which odor is used as reference: if C_z_ of 2H and C_y_ of EB are both isointense with C_x_ of ET, then C_z_ and C_y_ should also be isointense with each other (**Figure 3A**). For each odor pair, we held the concentrations of one odor (*reference* odor) constant, and varied the concentrations of the other odor. As above, we computed the difference between decision boundaries Δ_DB_ and the difference between the center of concentration ranges used in the session, ΔlogC. We used a used a linear fit to represent the relationship between Δ_DB_ and ΔlogC, and we estimated the PSE as the point for which Δ_DB_ = 0, as outlined above (Figure 3B, 3C, **3D**, see Methods). Isointense concentrations for ethyl tiglate, ethyl butyrate, 2-heptanone and acetophenone are reported in **Figure 3E**. To test for transitivity, we then compared, for each concentration range (that we call here low, medium or high) the concentration of 2H that was estimated isointense to EtT, to the concentration of EtB that was also estimated isointense to EtT. (**Figure 3A**). We found that there was no significant difference between the two estimations of 2H (**Figure 3E**). Transitivity was consistent at three intensity ranges. We conclude that the 2OCC paradigm provides a robust measure of isointense concentrations, and we leveraged it to compute isointense concentrations of four odorants (for four odorants: ethyl tiglate, ethyl butyrate, 2-heptanone, and acetophenone) for the three concentration ranges.

**Figure 3.**
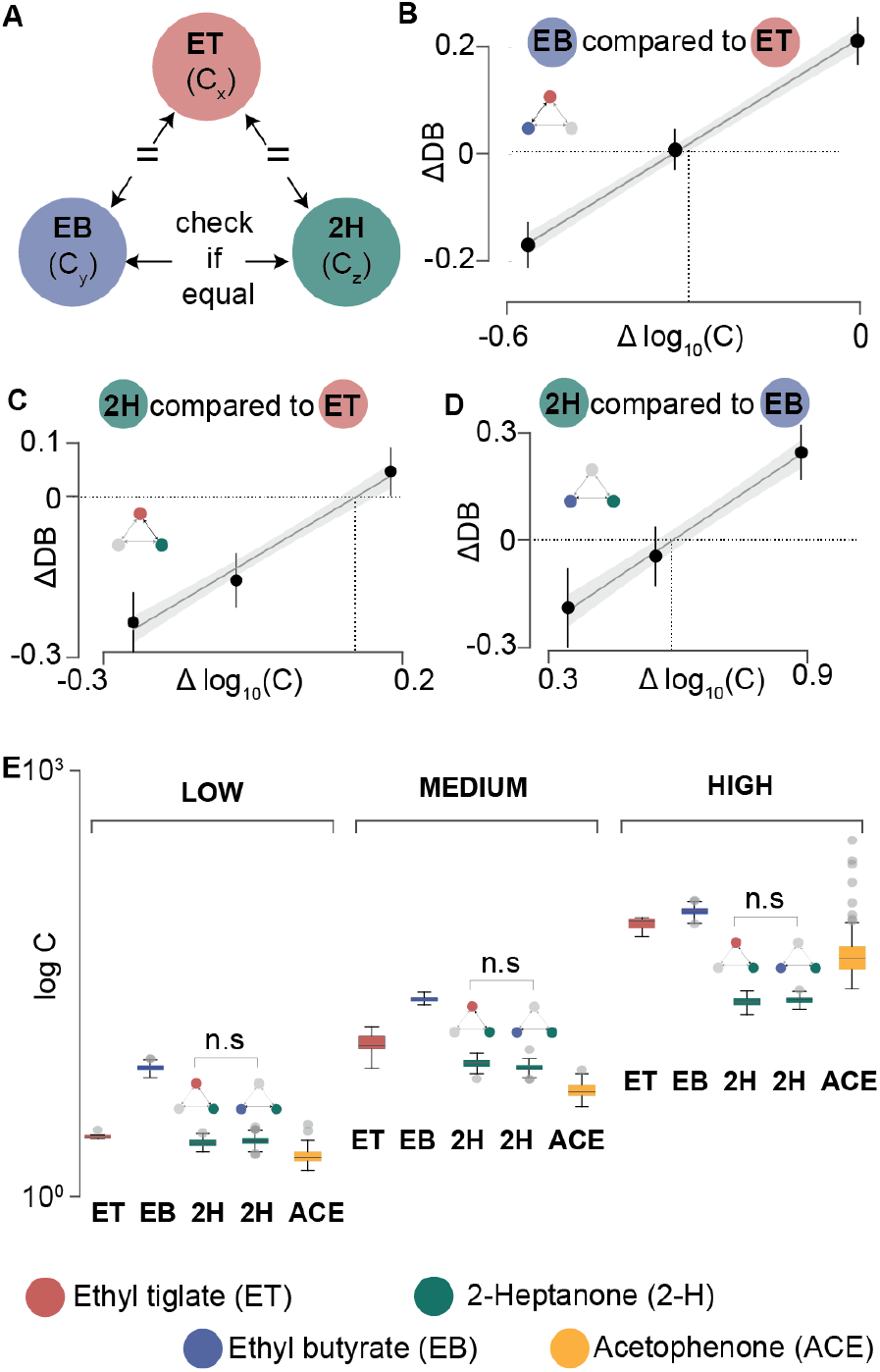
Isointensity is transitive in mice, enabling matched concentration estimates across odorants. **A**. Schematic illustrating the expected transitive property of intensity-matched concentrations. **B-D**. Performance mismatch index (Δ_DB_) as a function of concentration range differences for three odor pairs: (**B**) ET-EB, (**C**) ET-2H, and (**D)** EB-2H. Solid lines show linear fits across mice. Vertical dotted lines show the projection of Δ_ΔB_ = 0 from the regression line to the Δ*logx* axis. **E**. Intensity-matched concentration estimates for three odorants at low, medium, and high reference concentrations. Intensity-matched concentrations of 2H were estimated twice—once using ET as reference, and once using EB—to test transitivity.

## Discussion

This study characterizes odor intensity perception in a mouse model. Previous work used the 2OCC task to demonstrate that rats use a common perceptual scale to compare concentrations of different odors (Wojcik and Sirotin 2014). Here, we demonstrate that both mice and humans also evaluate concentrations of different odors using a common perceptual scale. Using explicit intensity ratings in humans, we demonstrate that this scale corresponds to perceived intensity. Applying the 2OCC paradigm in mice, we measured isointense concentrations for four odorants at three different intensities. This framework provides a foundation for a broader and more generalizable understanding of intensity perception.

### Measuring isointense odorant concentrations in mice

The ability to deliver controlled odor intensities in animal models opens multiple avenues in olfaction research. First, delivering stimuli at unknown intensities introduces unexplained variability into experiments aimed at understanding neural coding mechanisms. Odorants presented at mismatched intensities can evoke drastically different neural responses, but whether this is due to the intensity mismatch or actual difference in coding between the different odorants is unclear. Previous studies have attempted to disentangle which part of the neural code is relevant for odor quality perception by delivering a range of odorant concentrations and looking for parts of the neural code that are concentration-invariant (Bolding and Franks 2018; Cleland et al. 2012; Stern et al. 2018). However, increases in odorant concentration can also alter odorant quality perception (Gross-Isseroff and Lancet 1988; Laing et al. 2003; Conway et al. 2024), making this approach difficult to interpret. On the other hand, using isointense odorant stimuli would constitute an efficient and robust approach to investigate the neural code of odorant perception.

Second, delivering individual odorants at precise intensities is necessary for understanding the perception of odor mixtures. Natural olfactory stimuli are typically complex blends, and accurate reproduction of mixtures is necessary to generate more ethological olfactory stimuli, for example to study memory and navigation, or mating and parental behaviors (McRae et al. 2023; Marlin and Froemke 2017). A binary mixture of odorants A and B can be perceived as predominantly A, predominantly B, or a combination of both. Without knowing the relative perceived intensities of the components, predicting mixture perception is impossible.

Third, the neural representation of odor identity may shift across intensity regimes, as population responses shift from sparse to dense coding. Currently, it is unclear if experimental studies are performed at concentrations that are relevant and salient for animals in their natural environment (Wachowiak et al. 2025). Identifying ethologically relevant intensity ranges for normal olfactory function will allows us to better study the neural code of the olfactory system. In summary, to elevate olfactory stimulus control to the level of vision and audition, we must first establish reliable measures of perceived odor intensity.

### Similar psychophysical mechanisms of intensity perception across mice and humans

Our findings revealed that mice and humans use similar psychophysical mechanisms to evaluate odorant concentrations, applying a common perceptual scale across odorants. This suggests that stimulus amplitude is encoded along a unified internal axis of perceived intensity that enables cross-odorant comparison. In humans, this is consistent with psychophysical literature showing that the relationship between concentration and intensity is odorant specific. The parallel between mice and humans supports the idea that olfactory systems, despite anatomical and ecological differences, may share conserved computational strategies for extracting behaviorally relevant information from stimulus magnitude. This supports the use of mice as a model system for investigating the neural encoding of intensity perception and for translating insights into human sensory function.

### Understanding the neural encoding of intensity to improve human health

Multiple hypotheses of odor intensity coding have emerged from studies examining neural responses to changes in odorant concentration at several different levels of the olfactory system: firing rates (Wachowiak and Cohen 2001; Rospars et al. 2000; Duchamp-Viret et al. 1999; Bathellier et al. 2008), population size (Wachowiak and Cohen 2001; Bathellier et al. 2008; Rospars et al. 2008; Ma and Shepherd 2000) and activation latencies (Duchamp-Viret et al. 1999; Spors et al. 2006; Bolding and Franks 2017; Cang and Isaacson 2003) have all been shown to correlate with concentration. However, odor concentration is often a poor proxy for perceived intensity. At the same concentration, some odors evoke strong sensations, while others are weak or undetectable. Identifying isointense concentrations of different odors allows for more stringent experiments, as a neural code for odor intensity should both encode perceived intensity as stimulus concentration is altered and remain constant across different stimuli that have the same perceived intensity.

This approach can also be generalized to other sensory systems. The 2OCC task could be adapted to identify isointense tones in audition or iso-bright colors in vision. Comparative studies of intensity perception across sensory systems would provide a deeper understanding of how the brain encodes stimulus magnitude. This work has direct clinical relevance: atypical sensory intensity processing, such as hypo- and hyper sensitivities, is common across neurological and psychiatric conditions including autism spectrum disorder (Ashwin et al. 2014), and post-traumatic stress disorder (Engel-Yeger et al. 2013). These symptoms, which affect up to 16% of the general population (Yochman et al. 2013), can severely impair quality of life by disrupting engagement with the surrounding environment, leading to social withdrawal, isolation, or impaired work and daily activities. Yet the neural underpinnings of altered intensity perception remain poorly understood. Which brain areas are responsible for altered sensory encoding?

Which neural firing patterns are altered? This knowledge gap greatly complicates designing targeted therapeutic solutions for different symptoms. By characterizing odor intensity perception in the mouse this work establishes a platform for probing the neural basis of intensity coding at a mechanistic level. These insights will be critical for developing therapeutic strategies and preventative approaches for sensory disorders linked to altered intensity perception.

## Acknowledgements

This research was supported in part by grants from the NIH (U19NS112953, R01DC017757, R25 DC021651) and a Simons Foundation grant to BB (Simons Junior Fellowship 267385)

## Methods

### Mouse lines

All animal procedures were approved under a New York University Langone Health institutional animal care and use committee (IACUC) protocol 161211-02. Female C57BL/6J mice (Stock No: 000664, Jackson Laboratories) between 2 and 12 months old were used in all experiments and handled in accordance with institutional guidelines.

### Surgical procedures

Mice were anesthetized with isoflurane (2.0% during induction, 1.5% during surgery) during all surgical implantations. The skin covering the skull was removed and a custom 3D-printed headpost was fixed to the skull using dental cement (C&B Metabond, Parkell). Each animal recovered for at least 10 days prior to experiments.

### Odorant preparation and concentration calibration

Ethyl tiglate (CAS# 5837-78-5), Ethyl Butyrate (CAS# **105-54-4**) and 2-Heptanone (CAS#110-43-0) were purchased from Sigma Aldrich. The maximum concentration delivered in each session was determined by diluting the odorant in mineral oil. The air-phase concentration in each vial was then diluted through an air dilution olfactometer to obtain eight concentrations ranging from 0.01 to 0.1 the vial air-phase concentration. The air-diluted concentrations were equally spaced on a logarithmic axis. Air dilution was achieved through a custom-designed olfactometer (modified from Nakayama). Each olfactometer comprised a mass flow controller (MFC) (Alicat MC 1SLPMD/ 5M/5IN) that maintained the total clean air flow at 1L/min, and an MFC that conveyed a modulable airflow through an odorant vial (0–100 mL/min, Alicat, MC-100SCCMD/5M/5IN). The olfactometer also included an inline Teflon four-valve manifold, (NReserach, 225T082), one on-off three-port bypass valve (NResearch, TI1403270), and four odor vials. The headspace concentration of each vial expressed in parts-per-million was computed through a calibration procedure. In this procedure, we used a photoionization detector (200B: miniPID Fast Response Olfaction Sensor, Aurora, ON, Canada) to measure the concentration of each vial(Jennings et al. 2023). The PID outputs a voltage signal that is linearly proportional to the concentration of volatile organic compounds (VOCs) in the surrounding air.

The sensitivity of the PID is odorant-dependent, and PID signals do not directly represent the absolute concentration of VOCs. Therefore, to estimate the part-per-million concentrations of a vial headspace we compared the PID output from that vial to the PID output from a vial containing pure odor. We computed the ppm concentrations of the pure odorant as

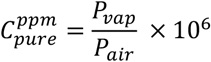

Where *P*_*vap*_ is the saturated vapor pressure of the odorant and *P*_*air*_ is the pressure of the air in contact with the odorant, which is ∼ 1atm (atmospheric pressure). We then derived the concentration of vials containing liquid dilutioins from the PID voltage signals as

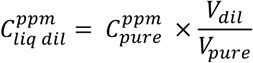

Where *V*_*dil*_ and *V*_*pure*_ are the steady state values of the PID voltage signals corresponding to the liquid dilution and pure odorant respectively, and 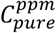 is the concentration of a vial containing pure odorant expressed in ppm. This procedure was repeated daily for all odorant vials used during behavior.

### Mice behavioral experiments

All behavioral experiments were controlled by a custom code software suite controlling a Bpod behavioral board (Sanworks, Bpod State Machine r2+). The software suite was written in Python (https://github.com/olfa-lab/PyBpodGUI). Mice were water deprived (1 ml/day) and habituated to handling for one week prior to the start of the experiments. The weight of the animal was monitored daily. Since mice were housed on a reverse light/dark cycle, training took place during the day. For 1-2 days mice were habituated to a head fixation system that left them free to run on a wheel. Mice received an additional 0.5 ml of water in the head fixation system during this part of the training. Animals were then trained to receive small droplets (∼3 μ l) of water by alternatively licking two small tubes. The tubes were symmetrically placed on the left and right of his mouth, at a comfortable distance for the animal to lick. Licks were detected through a capacitive touch sensor (Sparkfun) that was soldered to the tube. Water reached the tube through a hypodermic tube. When the animal licked the tubes water release was triggered through the closed-loop control of a pinch valve. When the animals learned to reliably lick for water and were able to receive their daily water dose in the experimental setup, they were trained to perform a two-choice task. Each trial was composed by a variable pre-stimulus period (∼3 s), a stimulus period (0.8 s) and a response period (5s) followed by a variable inter-trial-interval with variable duration (6-12 s). Correct licking was rewarded with a 5 ul water drop, while incorrect licking resulted in a 1 second white-noise tone. Outside of the response window animals were left free to lick, but no water was delivered in response to licking activity. The respiration signal was monitored through a pressure sensor coupled to a Teflon odor port that passed air at a steady rate of 0.5 l/min (Wilson et al. 2017). The stimulus was delivered upon detection of the animals’ exhalation, to guarantee odor presence by the time of the animal inhalation. Mice were initially presented with either a low or high concentration of the odorant ethyl tiglate (*reference* odor). Target concentrations were achieved through a combination of liquid and air dilution (see Odorant preparation and concentration calibration). The low concentration was equivalent to one tenth of the high concentration, as expressed in part per millions. Mice were rewarded to lick the left tube in response to the low concentration, and the right tube in response to the higher concentration. When animals reached a 75% performance, two more concentrations within the were added to the task. The additional concentrations were comprised between the low and high concentrations initially used and equally spaced on a logarithmic axis. When mice reached proficiency with the four stimuli paradigm, four additional concentrations were added to the task, for a total of eight stimuli. The progression through the different training phase was personalized to each animal depending on their level of proficiency. The animals were rewarded for licking left for the four lowest concentrations, and for licking the right tube for the four highest concentrations. The training phase ended when mice reached a performance > 75% and the decision boundary was centered in the tested concentrations range and stable across session (**Figure S1D**). We computed the decision boundary as the concentration associated to equal probability of licking right or left (see Human intensity ratings

### curves

Hill parameters were extracted by fitting a Hill function to concentration-intensity data points. Datapoints from all subjects were combined in one single dataset. Noise was estimated through bootstrapping (sampling with replacement, N = 1000). Each bootstrapped dataset was used to fit a Hill curve for each odorant ***o***:

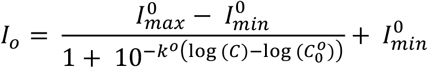

where *I*^*o*^_*max*_, *I*^*o*^*min, j*^*o*^ and *C*^*o*^_*0*_ are odor specific parameters. Intuitively, *I*_*max*_ is the maximal relative intensity value reached by that odorant, *I*_*max*_ is the minimum relative intensity value reached by that odorant, *j* determines the steepness of intensity increases with concentration, and *C*_*0*_ corresponds to the inflection point of the curve. During the fitting procedure, *Io*_*max*_ and *Io*_*min*_ were constrained in the [0, 100] range, *j*^*o*^ in the [10-6, 5] range and *Co0* in the [-8, -1] range on a logarithmic scale.

### Behavioral data analysis

At the end of the training phase, we repeated the same paradigm with a second odor, that was novel to the mouse. The concentrations of this second odor were selected arbitrarily at a similar concentration range that the mouse was trained on, but without guaranteeing perfect concentration matches across odors. Mice were able to perform the task for the novel odor and achieved performances > 75% within a couple of sessions. Once checked that mice were able to generalize to different odorants, we combined two odorants in a single experimental session. We alternatively delivered one of eight concentrations of both odorants, for a total number of 16 different stimuli. For each odorant, mice were rewarded for licking left the four lowest stimuli and for licking right for the four highest concentrations. We only run one session per day, for a total number of 16 different stimuli per session. Experimental sessions were terminated when animals did not respond to the odor stimuli for ten consecutive trials. The concentration of the first odor the mice were introduced to (or *reference* odor) was always kept fixed at the original value, while the concentration of the second odor (*measured* odor) was increased and decreases. The concentration values of the measured odor were selected daily based on the results of previous experimental days, in order to equally represent sample concentration pairs that yield positive, negative intensity matching indexes. The maximal concentration for each odor was always obtained by liquid dilution. Each liquid dilutions was appropriately calibrated to estimate the absolute parts per million value (**Figure S1B**, see Odorant preparation and concentration calibration).

### Human behavioral experiments

#### Participants

Seventeen individuals (9 females, 8 males) with ages ranging between 19 and 50 (mean = 32.1, SD = 10.54) participated in the study. All participants reported a normal sense of smell and were recruited from the Philadelphia area. In addition, participants reported no health complications related to a diminished sense of smell such as renal disease, neurodegenerative disease, or congestion at the time of testing. The research protocol was approved by the University of Pennsylvania IRB (#818208), and all participants gave informed consent prior to enrolling in the study. All experiments were conducted in rooms specially constructed for human olfaction experiments at the Monell Chemical Senses Center. The rooms are well-ventilated and painted minimally in off-white. During the testing portion of session, the experimenter, olfactometer and participant were visually separated. All experimental measures were done to minimize cross-trial contaminations and panelist bias.

### Stimuli

Acetal (CAS#105-57-7) and 2-Heptanone (CAS#110-43-0) were used as odorant stimuli. Both odorants were purchased from Sigma-Alpha and delivered via a controlled odor delivery system using gas-sampling bags (Pellegrino et al. 2025). This approach ensures consistent odorant concentrations throughout testing sessions and improves measurement reliability compared to conventional methods. We constructed an odor delivery system using Nalophan gas sampling bags (Miller and McGinley 2008) sealed at both ends with cable ties: one end around a tube fitted with an external septum, the other around an open/close valve. Bags were first evacuated via vacuum through the valve, then filled with dehumidified, carbon-filtered air at a flow rate of 2 LPM for 5 minutes to achieve a volume of 10 L. The liquid odorant was injected through the septum and allowed to equilibrate into the gas phase, a process that took between minutes to hours depending on the odorant’s volume and volatility. After reaching equilibrium, the odorant concentration was uniform throughout the bag. For sampling, we connected the open/close valve to a flexible nose mask equipped with a low-resistance one-way valve. This system allowed natural inhalation of the headspace while preventing ambient air contamination. The mask’s design enabled a tight facial seal and quick attachment to multiple bags in succession.

### Experimental Procedure

The study consisted of seven sessions spaced one week apart. Session 1 collected concentration-intensity functions for the odorants through an explicit intensity rating task, while Sessions 2-4 employed these data to design matched intensity tasks. For the explicit intensity rating task (Session 1), subjects were trained to rate intensity with the general Labeled Magnitude Scale (Green et al. 1996; Bartoshuk et al. 2003) (gLMS) using examples from other modalities (e.g. “please rate the loudness of a jet engine”) and were asked to disregard pleasantness when rating intensity. Additionally, participants rated the intensity of blinded weights. After training, panelists were trained on odor sampling with the olfactometer with a few random low and high intense odorants. This practice was necessary to get the participants to time their sniff to the delivery of the odorant (which was cued via a countdown; “3”, “2”, “1”, “smell now”). Odorants were sampled at up to eight concentrations in duplicate to obtain a proper concentration-intensity curves. The concentrations used can be found in Supplementary Table 1. A duplicate rating of a blank (clean air) were collected to obtain a measure of baseline. All stimuli were randomized across panelists with an interstimulus interval of 25s.

In Sessions 2-7 participants performed the 2OCC task. Acetal was used as a *reference* odor (concentration was kept constant) and 2-Heptanone as the *measured* odor (concentration was shifted session by session). This procedure was repeated at two concentration ranges, one centered around the concentration that produced an intensity percept equal to 15% (EC15) to the maximal perceived intensity of 2-heptanone and one centered around the concentration that produced an intensity percept equal to 50% (EC50) to the maximal perceived intensity of 2-heptanone. At the EC15 range, the measured odor concentration was shifted (above and below) of 0.25 on the logarithmic scale. At the EC50 range, the measured odor concentration was shifted of 0.5 on the logarithmic scale. For each concentration range, in the first behavioral session, participants were presented with isointense concentrations, in the second behavioral session, the concentration of 2-heptanone was increased and in the third behavioral session the concentration of 2-heptanone was decreased. Before the matched intensity tasks, participants were trained to classify stimuli as “Low” or “High” intensity. Training involved the highest and lowest intensities of both odorants to familiarize participants with the task. During both training and testing, stimuli were delivered following the same countdown, and participants classified each as “Low” or “High,” receiving feedback on accuracy. Each stimulus was presented four times in random order, with a 25-second interstimulus interval.

The intensity-matched concentrations in used in the first behavioral sessions were derived from the concentration-intensity functions, modeled as a Hill function, from the intensity ratings (see Data Analysis, Human intensity ratings curves).

## Data analysis

### Human intensity ratings curves

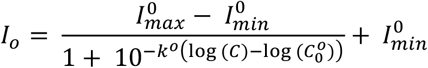

where *I*^*o*^_*max*_, *I*^*o*^_*min*_, *j*^*o*^ and *c*^*o*^_*0*_ are odor specific parameters. Intuitively, *I*_*max*_is the maximal relative intensity value reached by that odorant, *I*_*max*_is the minimum relative intensity value reached by that odorant, *j* determines the steepness of intensity increases with concentration, and cocorresponds to the inflection point of the curve. During the fitting procedure, *I*^*o*^_*max*_ and *I*^*o*^_*max*_ were constrained in the [0, 100] range, *j*^*o*^ in the [10^−6^, 5] range and *C*^*o*^_*0*_ in the [-8, -1] range on a logarithmic scale.

### Behavioral data analysis

For both mice and humans, behavioral raw data consisted in binary classifications (0 or 1) of different odorant concentrations. To estimate the variability in the behavioral data, we implemented a bootstrap procedure on the raw behavioral data. We used 100 iterations and, in each iteration, random samples of the same size as the original dataset were drawn with replacement. Discrete performance values for each concentration were computed as the percentage of “high-like” responses. The performance statistics were estimated by computing mean, standard deviation and standard error of the mean of the binary vector of all responses at each concentration. To estimate a continuous performance curve, raw data were used to fit a sigmoidal function expressed as 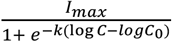, with *I*_*max*_, *C*_*o*_ and *k* as free parameters. To estimate the decision boundary for each performance curve, we computed the point at which the sigmoid crossed the 50% performance value. We normalized those values to fall in a normalized logarithmic scale ranging between 1 and 2. Those values represent 10% and 100% of the vial saturated headspace, that we diluted through an air dilution olfactometer. Decision boundaries were then used to estimate the relative intensity mismatch index (Δ_DB_). The relative intensity mismatch index was computed as the difference between the decision boundary of the reference odor and the measured odor, on the normalized logarithmic concentration axis. This resulted in positive a Δ*I* value associated to increased rates of “high-like” responses for the measured odor. The difference in concentrations associated to each Δ*I* was computed as the difference between the concentration of the measured odor and the reference odor on the logarithmic axis (Δ*log*C = *log*C_meas_–*log*C_ref_). We performed a linear regression between intensity mismatch index (Δ_DB_) and the concentration difference (Δ*log*C). We used this regression to compute the value of (Δ*log*C that yielded a value of Δ*I* = 0. We estimated isointense concentration of the measured odor with respect to the reference odor as:

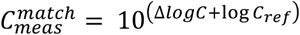

In the mouse, we repeated this procedure with the following odor-pairs: ethyl tiglate and ethyl butyrate, ethyl tiglate and 2-heptanone, ethyl butyrate and 2-heptanone, ethyltiglate and acetophenone. For the ethyl butyrate/2-heptanone pair, the reference concentration of ethyl butyrate was chosen to be isointense to the reference concentration of ethyltiglate. This additional step was added to verify that the estimation of isointense concentrations was not dependent on the specific odor-pair selected.

In human participant, we performed this procedure for the 2-heptanone and acetal odor pair.

### Statistic

In Figure 1B, we compared performance across conditions (ethyl tiglate, ethyl butyrate and 2-heptanone), using the Friedman test, (level of significance ^*^ p <0.05).

In Figure 2E, 3F, statistics and significance were evaluated by estimating two-sided residuals via bootstrapping (^*^ p < 0.05).

The p-value was estimated as:

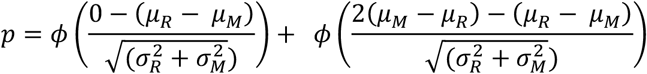

Where *Φ* is the cumulative distribution function (CDF) of the standard normal distribution, *μ*_*R*_ and *μ*_*M*_ are the means of the two bootstrapped distributions R and M and *σ*_*R*_ and *σ*_*M*_ are the standard deviations of the two bootstrapped distributions. The formula uses the cumulative distribution function (CDF) of the normal distribution to calculate the probability that the difference between the means of the two distributions *μ*_*R*_ − *μ*_*M*_, is consistent with a null hypothesis that assumes no significant difference between them. The term 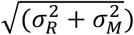represents the combined variance of the two distributions. The first term:

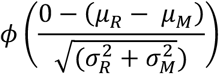

calculates the probability of the observed difference being less than or equal to zero, while the second term,

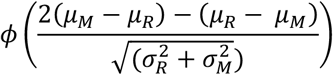

accounts for the tail on the other side of the distribution. Together, these components provide the overall p-value, which quantifies the likelihood of observing a difference between the means of the two distributions purely by chance.

In Figure S1, a one-sample t-test was applied despite the small sample size (n = 4), under the assumption of approximate normality, supported by the lack of gross deviations in data distribution and the known properties of the underlying measurement (Shapiro test, p> 0.05). All other statistical tests were performed with the Mann-Whitney U test, levels of significance (* p <0.05, ^**^ p < 0.01, ^***^ p < 0.001)

---

**Supplementary Table 1:** Concentrations of 2-heptanone and acetal used for intensity rating task.

| Odorant | Concentration (v/v) |  |  |  |  |  |  |  |
| --- | --- | --- | --- | --- | --- | --- | --- | --- |
| Index | 1 | 2 | 3 | 4 | 5 | 6 | 7 | 8 |
| 2-Heptanone | -7.6 | -7. | -6.5 | -5.82 | -5.30 | -4.69 | -4.22 | -3.88 |
| Acetal | -7.69 | -6.52 | -5.5 | -4.82 | -4.12 | -3.64 | -3.49 |  |

