## Supplementary Information for "Mice and humans evaluate odor stimulus strength using common psychophysical principles"

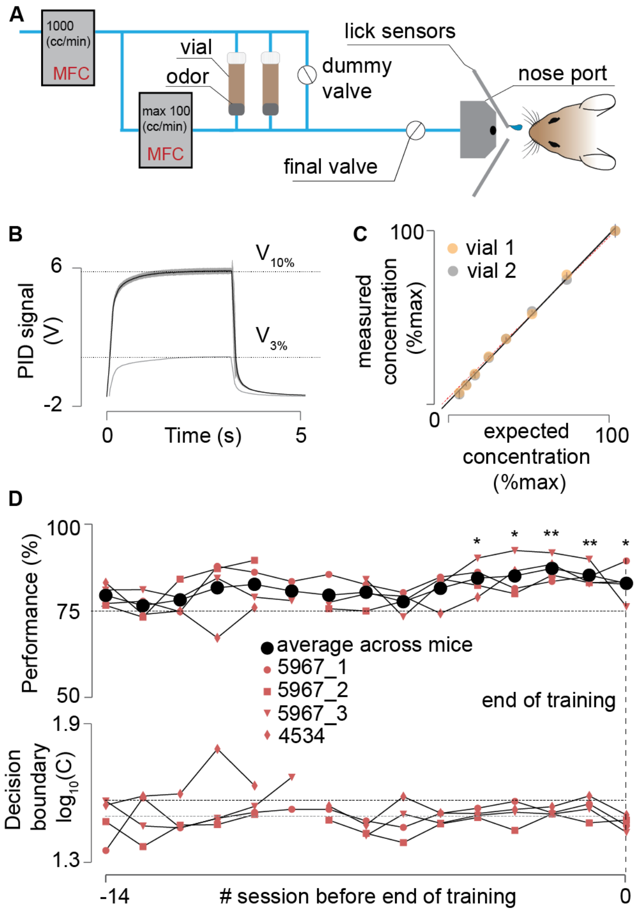


**Figure S1. Experimental setup and task training. A.** Schematic of the air-dilution olfactometer. Two mass-flow controllers set the ratio of odorized vs clean air delivered to the mouse. The mouse inserted its nose into a port, and lick responses were recorded by two capacitive lick sensors, which also delivered water rewards. **B.** Example odor delivery traces recorded with a photo-ionization detector (PID). To estimate the effective air concentration obtained by diluting odorants in a solvent, PID traces from different liquid dilutions were compared to traces from a 10% air dilution of a pure odorant vial. **C.** Relationship between imposed air-dilution factors and the effective air-dilution factor, measured as the ratios between saturation values of PID traces at different air-dilution factors. Dots show averages; error bars show standard deviation. **D.** Evolution of average performance (top) and decision boundary (bottom) for each mouse over the last 14 training sessions. Training was terminated once mice reached ≥ 75% performance (one-sample t-test, $p<0.05$, under the assumption of approximate normality from Shapiro test $p>0.05$) and maintained a stable decision boundary within ±0.071 log units of the midpoint of the concentration range for at least three consecutive days.

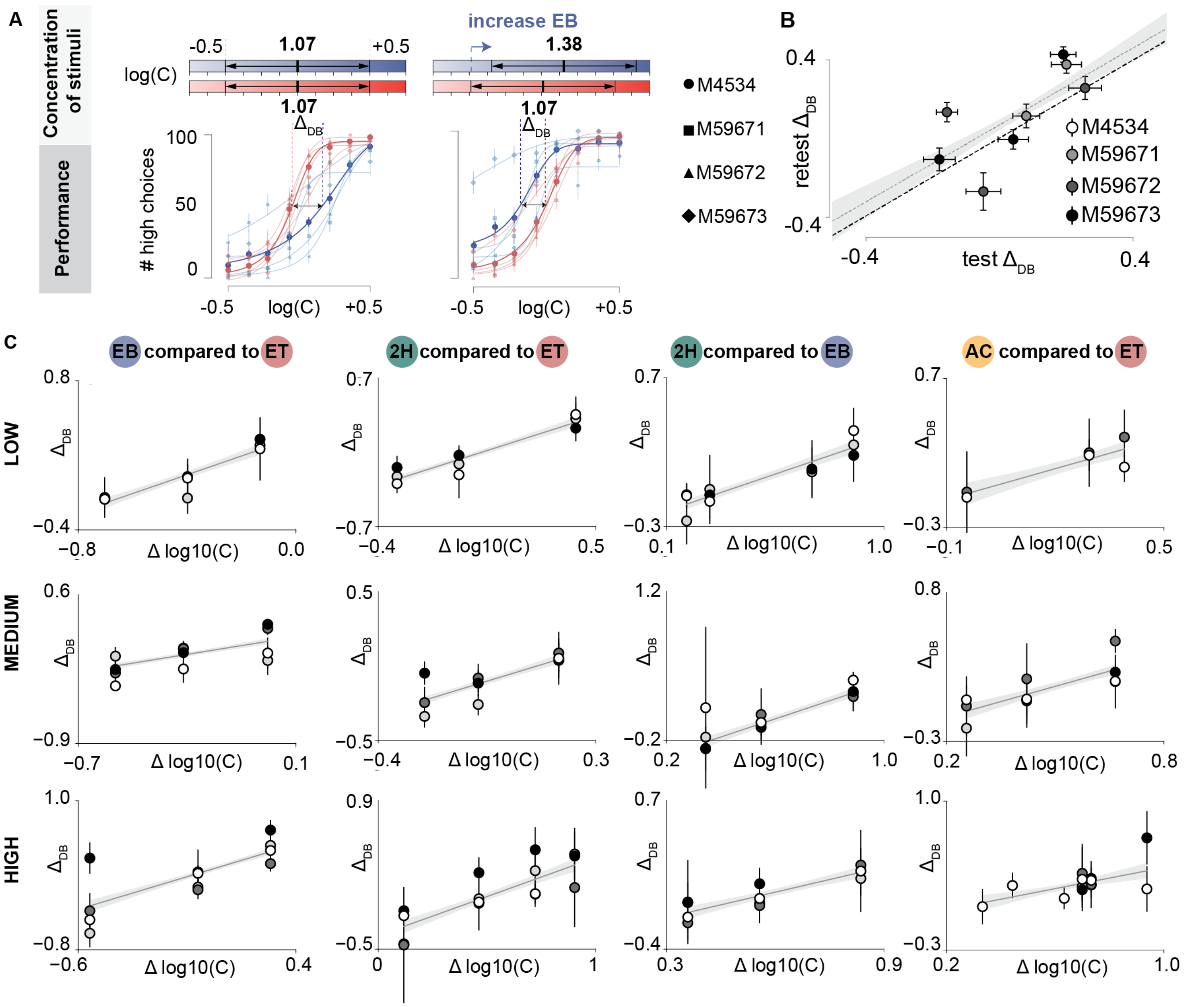


**Figure S2. Isointense concentrations for four odorants across three intensity ranges. A.** Individual performance curves of n=4 mice (transparent lines and dots) overlaid with the mean performance across animals during the two-odorant task with ethyl tiglate (red) and ethyl butyrate (blue). Data correspond to the concentration range combinations shown in Figure 1E-F. **B.** Test-retest values of $\Delta_{DB}$ for each animal at the medium intensity range. Error bars indicate SEM of estimated $\Delta_{DB}$ for each session (test or retest). The gray dotted line shows the linear fit across all test-retest data points, with the shaded region indicating the standard deviation of the fits. The black dotted line represents the unity line for reference. **C.** Performance mismatch index ($\Delta_{DB}$) as a function of concentration range differences for four odor pairs (columns) at three intensity ranges (low, medium and high; rows). Each data point represents the session average $\Delta_{DB}$ of one animal. Error bars show SEM per session. Solid lines show linear fits across mice, with shaded areas denoting the SEM of the fit.
